# WNT responsive SUMOylation of ZIC5 exerts multiple effects on transcription to promote murine neural crest cell development

**DOI:** 10.1101/2020.11.04.369124

**Authors:** Radiya G. Ali, Helen M. Bellchambers, Nicholas Warr, Jehangir N. Ahmed, Kristen S. Barratt, Kieran Neill, Koula E. M. Diamand, Ruth M. Arkell

## Abstract

Zinc finger of the cerebellum (Zic) proteins act as classical transcription factors to promote transcription of the *Foxd3* gene during neural crest cell specification. Additionally, they can act as co-factors that bind TCF molecules to repress WNT/β-catenin dependent transcription without contacting DNA. Here, we show ZIC activity at the neural plate border is influenced by WNT-dependent SUMOylation. In a high WNT environment, a lysine within the highly conserved ZF-NC domain of ZIC5 is SUMOylated, which decreases formation of the TCF/ZIC co-repressor complex and shifts the balance towards transcription factor function. The modification is critical *in vivo*, as a ZIC5 SUMO-incompetent mouse strain exhibits neural crest specification defects. This work reveals the function of the ZIC ZF-NC domain, provides *in vivo* validation of target protein SUMOylation, and demonstrates that WNT/β-catenin signalling directs transcription at non-TCF DNA binding sites. Furthermore, it can explain how WNT signals convert a broad domain of *Zic* ectodermal expression into a restricted domain of neural crest cell specification.

## Introduction

The neural crest is a transitory population of multi-potent cells that arises during gastrulation. Inductive signals from various growth factors, including WNTs, establish the neural plate border (NPB) at the juncture of the neural and non-neural ectoderm. The NPB contains the prospective neural crest and expresses multiple transcription factors, including members of the *Zic* gene family (Stuhlmiller and García-Castro, 2012; Bronner and Simões-Costa, 2016; Rogers and Nie, 2018; York and McCauley, 2020). These transcription factors, in response to sustained WNT signals, cooperate to direct the expression of a second set of transcription factors, known as neural crest specifier genes, including *Foxd3*. In chick embryos, ZIC1 binds to and promotes expression from a conserved enhancer element containing a Zic responsive element (ZRE) upstream of the *FOXD3* gene (Simões-Costa *et al.*, 2012). In mouse embryos, *Zic1* is not expressed during neural crest cell (NCC) induction (Elms *et al.*, 2004), but three closely related genes are: *Zic2*, *Zic3* and *Zic5* (Furushima *et al.*, 2000; Elms *et al.*, 2004). Mouse embryos that lack either functioning ZIC2 or ZIC5 protein have depleted neural crest production (Elms *et al.*, 2003; Inoue *et al.*, 2004); thus these genes likely encode putative endogenous murine *Foxd3* expression regulators.

ZIC proteins can also act as co-factors. For example, Zic family members have been shown to bind TCF proteins (Pourebrahim *et al.*, 2011; Fujimi, Hatayama and Aruga, 2012; Zhao *et al.*, 2019) and inhibit β-catenin/TCF mediated transcription at WNT responsive elements (WREs) stimulated by canonical WNT signalling (Pourebrahim *et al.*, 2011; Fujimi, Hatayama and Aruga, 2012). In a low WNT environment (i.e. when β-catenin protein is degraded by the β-catenin destruction complex), WREs are often repressed by transcriptional mediators from the TCF/LEF family in concert with co-repressors, the best studied of which are members of the TLE family of Groucho related co-repressors (Ramakrishnan *et al.*, 2018). ZIC proteins can also interact with TCF proteins to function as co-repressors (without contacting the DNA at WREs) in cultured mammalian cells, and in *Xenopus* and zebrafish embryos (Pourebrahim *et al.*, 2011; Fujimi, Hatayama and Aruga, 2012; Zhao *et al.*, 2019). Upon WNT stimulation and nuclear import of β-catenin (as occurs at the NPB; Ferrer-Vaquer *et al.*, 2010), the repressor complex on a WRE is converted to a TCF/β-catenin activation complex (Gammons and Bienz, 2018). Exactly how the various molecular functions of ZIC proteins are controlled to ensure the timely activation of the neural crest specifier genes in a high WNT environment is unknown (Ali, Bellchambers and Arkell, 2012; Houtmeyers *et al.*, 2013).

One mechanism by which protein activities are dynamically regulated is via post translational modification (PTM). SUMOylation, in which the SUMO (Small Ubiquitin-like Modifier) protein is reversibly attached to specific lysine residues of target proteins, tends to alter the interaction of its target substrates with other proteins or DNA. It can do this by enhancing or blocking interaction sites or by inducing conformational change in the target protein (Henley, Craig and Wilkinson, 2014; Hendriks and Vertegaal, 2016; Han *et al.*, 2018). Additionally, once SUMOylated, the protein may be targeted by SUMO-targeted ubiquitin ligases that specifically recognise and ubiquitylate SUMOylated proteins. Thus, SUMO conjugation to a target protein can result in a range of functional changes (altered DNA binding, protein-protein interactions, subcellular localisation or protein stability) (Wei, Schöler and Atchison, 2007; Kim, Chia and Costantini, 2008; Choi *et al.*, 2011; Chen *et al.*, 2013). The SUMOylation cycle, in which SUMO is matured, activated and passed to the conjugating enzyme and then (in combination with an E3 ligase) is conjugated to a target lysine in a substrate protein, differs slightly from that of ubiquitylation. For example, though mammalian cells express at least three different SUMO protein isoforms (SUMO1-3), there is only one SUMO conjugating enzyme (UBC9) and it plays a role in target specificity by binding to a SUMOylation consensus site in proteins. However, this binding is relatively weak and SUMOylation of most substrates is inefficient in *in vitro* reactions lacking the appropriate E3 ligase (Jakobs *et al.*, 2007; Varejão *et al.*, 2020). *Ubc9* is expressed specifically at the NPB in chick embryos and its depletion downregulates the expression of NCC specifier transcription factors (Luan *et al.*, 2013), demonstrating that this PTM is essential for timely NCC development in the chick. Additionally, SUMOylation of the NPB transcription factor PAX7 in chick embryos (Luan *et al.*, 2013) and of the NCC specifier transcription factors Sox9 and Sox10 in *Xenopus* and chick embryos (Taylor and LaBonne, 2005; Liu *et al.*, 2013) influences NCC development. Little is known, however, about the role of SUMOylation in mammalian neural crest specification.

Here, we investigate SUMOylation of the multifunctional transcription regulator ZIC5 and explore the functional consequences of ZIC dependent SUMOylation in cells, at the murine NPB and in varying WNT environments. We report that ZIC5 is poly-SUMOylated at a deeply conserved lysine and that conservative substitution of this single lysine with a non-modifiable arginine residue disrupts NCC development during murine embryogenesis. Cell-based investigations of the functional consequence of ZIC5 SUMOylation demonstrate that SUMOylation decreases co-repressor activity and potentiates the trans-activation ability of ZIC5, including at the ZIC responsive element in a *Foxd3* enhancer. Moreover, we find that stimulation of the canonical WNT pathway increases the proportion of the ZIC5 protein that is SUMOylated and that, in this high WNT environment, ZIC5 co-repression of WREs is diminished. This SUMOylation driven, bi-phasic response of ZIC5 to WNT signalling can theoretically influence transcription at both WREs and ZREs. It also provides a conceptual basis to explain the fact that NCC specification occurs in one region within a broad neuroectodermal domain of ZIC expression: high concentration canonical WNT signals at the NPB convert ZIC and TCF proteins into transactivators, whereas repression at WREs persists in the future lateral neuroectoderm.

## Results

### ZIC5 is a target of the post-translational modification SUMOylation

To investigate the molecular mechanism regulating the balance between ZIC transcription factor and co-factor abilities, we focused on a small (14-21 aa in size), highly conserved, but functionally uncharacterised domain located immediately N-terminal of the zinc finger domain in each of the vertebrate Zic proteins, termed the Zinc Finger N-terminally Conserved domain (ZF-NC) (Aruga *et al.*, 2006). Analysis of the human ZF-NC region of ZIC1-5 identified a high probability consensus SUMOylation motif (Gareau and Lima, 2010) at the C-terminus of the domain (Figs. 1A and S1, Motif 1). This same motif was previously identified as a bona-fide SUMOylation site in the ZIC3 protein (Chen *et al.* 2013). Further investigation revealed that a subset of ZIC proteins (ZIC1, 3, 5) contain a second consensus SUMOylation motif located at the zinc finger 3 and 4 boundary (Fig. 1A and S1, Motif 2) and a third consensus close to the N-terminal of ZIC5 that is not conserved in other Zic family members (Fig. 1A, Motif 3).

**Figure 1:**
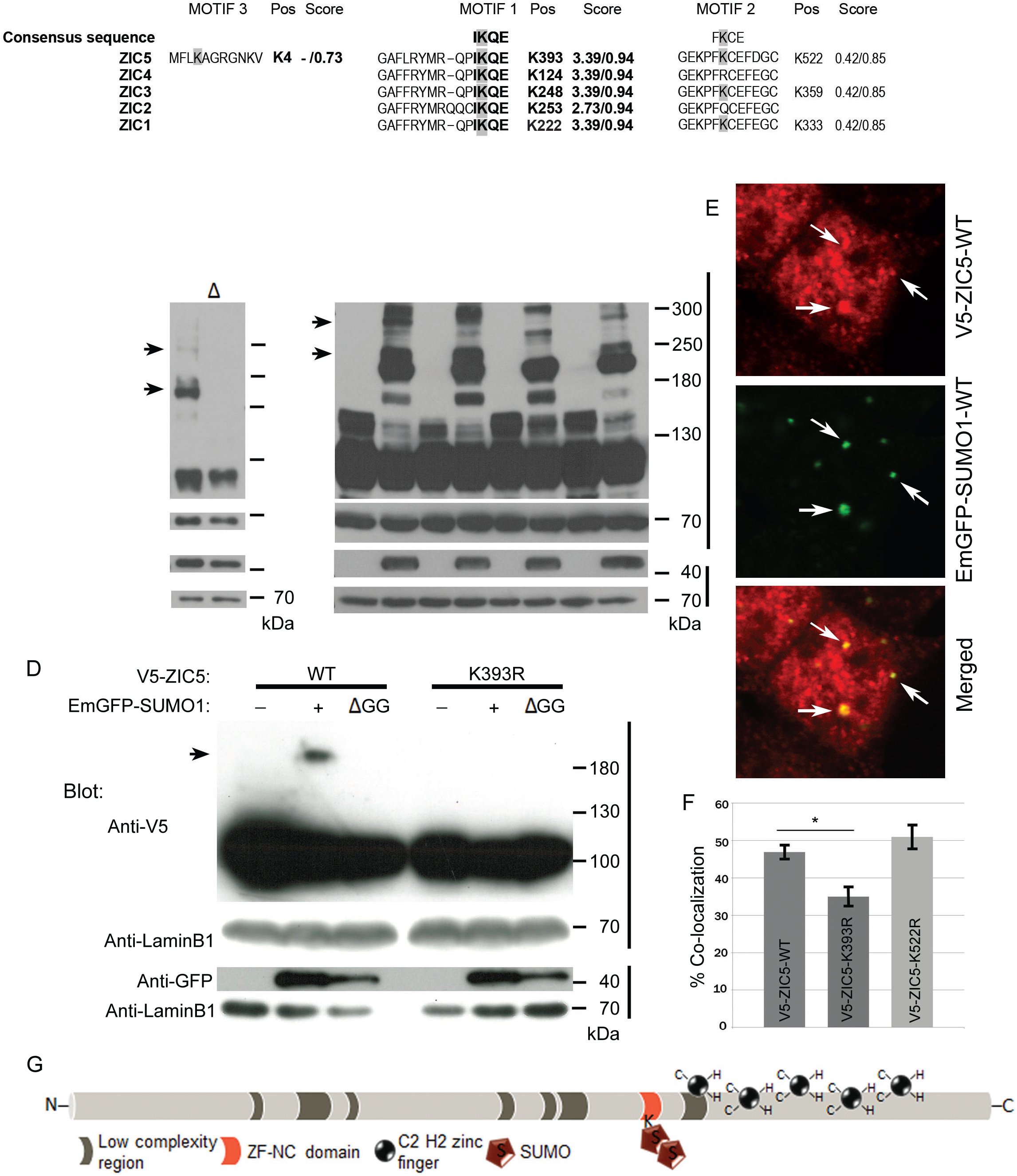
ZIC5 is SUMOylated at a conserved lysine within the ZF-NC domain. (A) Alignment of human ZIC proteins with putative SUMOylated lysines [K] highlighted. Motif 3 is not conserved in ZIC1-4. The first score for each Motif is computed by SUMOsp and the second by SUMOplot. (B,C) Representative WB of HEK293T cell nuclear fractions following transfection of V5-tagged UBC9-fused ZIC5 protein in the presence of the SUMO1 proteins indicated. (B) A series of higher MW products (arrows indicate bands of ~200 kDa and 280 kDa) are found in the presence of EmGFP-SUMO1-WT but not in the presence of the ΔGG inactive form. (C) Mutation of K393, but not K522 or K4, leads to depletion of the SUMO1 dependent higher MW products. Arrows indicate bands of ~200 kDa and 280 kDa and missing bands are indicated by asterisks. (D) WB of HEK293T cell nuclear fractions following transfection of the V5-tagged ZIC5 proteins in the presence of the SUMO1 proteins. One high molecular weight form of ZIC5 (arrow; ~200 kDa) is detected in the presence of WT SUMO1 but not in the presence of the ΔGG inactive form of SUMO1. The higher molecular form is not detected when K393 is mutated, indicating K393 is the sole target of ZIC5 SUMOylation. For B-D, n = 3 independent WBs, including loading control of the anti-V5 blot. WB to show overexpressed EmGFP-SUMO1 protein and corresponding loading control are at bottom of the panel. (E,F) Immunofluorescence analysis of ZIC5 and SUMO1 co-localisation of V5-ZIC5 (α-V5, red) and GFP-SUMO (α-GFP, green) in nuclear foci of transfected HEK293T cells. (E) Representative images, arrows indicate points of co-localisation. (F) Percentage of SUMO foci enriched for ZIC5. Pooled data from two independent experiments. Error bars = SE (regression analysis) *: *p*<0.001. (G) Schematic of ZIC5 showing location of SUMO attachment.

To determine whether the ZICs motifs are *bona fide* SUMOylation targets, we assessed whether one member of the ZIC family, ZIC5, can be SUMOylated by SUMO1 using the cell-based UBC9-fusion-directed SUMOylation system (UFDS) (Jakobs *et al.*, 2007). The direct fusion of UBC9 to the target protein catalyses SUMO E3 ligase-independent SUMOylation, avoiding the need for target specific endogenous ligases (Jakobs *et al.*, 2007). HEK293T cells transiently expressing V5 epitope-tagged wild-type human ZIC5 fused to UBC9 (V5-UBC9-ZIC5-WT) alone or with either EmGFP-tagged wild-type SUMO1 (EmGFP-SUMO1-WT) or a SUMOylation-defective SUMO1 mutant (EmGFP-SUMO1-ΔGG) (Kamitani, Nguyen and Yeh, 1997) were lysed and subjected to SDS-PAGE and western blotting (WB). When V5-UBC9-ZIC5-WT and EmGFP-SUMO1-WT were co-expressed, additional heavier (~200 and 280 kDa) molecular weight (MW) bands of V5-UBC9-ZIC5-WT (base MW of ~120 kDa) were detected (Fig. 1B). These bands were not detected in the absence of EmGFP-SUMO1-WT or the presence of EmGFP-SUMO1-ΔGG, suggesting V5-UBC9-ZIC5-WT can be SUMOylated (Fig. 1B,C). To corroborate this finding, the behaviour of transiently expressed V5 epitope-tagged wild-type ZIC5 (V5-ZIC5-WT) in the absence or presence of EmGFP-SUMO1-WT or EmGFP-SUMO-ΔGG in HEK293T cells was compared. As before, an additional heavier MW band (~180 kDa) of V5-ZIC5-WT (base MW of ~100 kDa) was only observed in EmGFP-SUMO1-WT expressing cells (Fig. 1D).

The increase in the MW of exogenously expressed V5-ZIC5-WT (by ~80 kDa; Fig. 1D) is consistent with the addition of two EmGFP-SUMO1-WT molecules and suggests that ZIC5 is either multi-mono-SUMOylated (single SUMO conjugated at multiple sites) or poly-SUMOylated (a chain of SUMO molecules at a site; Gocke, Yu and Kang, 2005). To clarify this and identify genuine SUMOylation motifs within ZIC5, V5-tagged ZIC5 mutant constructs with a lysine (K) to arginine (R) mutation in either Motif 1 (V5-ZIC5-K393R) or Motif 2 (V5-ZIC5-K522R) were expressed in the presence or absence of EmGFP-SUMO1-WT. No increased MW band of V5-ZIC5-K393R was observed when co-expressed with EmGFP-SUMO1-WT (Fig. 1D), whereas V5-ZIC5-K522R was no different to wild-type (data not shown), indicating that K393 is the sole site of SUMO attachment and that K393R prevents SUMOylation. Even when the UFDS system was employed, mutation of Motif 2 or Motif 3 showed no evidence of affecting the PTM of ZIC5 (Fig. 1C).

SUMOylated proteins are reported to co-localise with SUMO1 foci or nuclear bodies (NB) that are thought to be sites of active SUMOylation (Navascués *et al.*, 2007; de Cristofaro *et al.*, 2009; Lallemand-Breitenbach and de Thé, 2018); inhibition of SUMOylation may therefore be expected to decrease the extent of co-localisation. Thus, to further validate our WB results we measured the proportion of EmGFP-SUMO1-WT foci that are enriched for exogenously expressed wild-type or mutant ZIC5 (Fig. 1E,F) in the nucleus of HEK293T cells. The co-localisation of EmGFP-SUMO1-WT with V5-ZIC5-K393R (35.1%, S.E. 2.89%) showed a statistically significant decrease compared to V5-ZIC5-WT (46.9% S.E. 2.12%; *p*<0.001), whilst V5-ZIC5-K522R showed no significant difference from V5-ZIC5-WT (51%, S.E. 3.52%). Although significantly reduced, the K393R mutation did not ablate co-localisation with SUMO1. Extrapolating from the observations of Gocke *et. al.* (2005), we speculate that only a small fraction of the V5-ZIC5-WT in SUMO1 NB is SUMOylated at a given time and that localisation in these SUMO1 enriched zones might aid rapid SUMOylation of ZIC5 in response to cellular cues. Taken together, these data demonstrate that ZIC5 is poly-SUMOylated at a single SUMOylation motif (Motif 1, K393R) (Fig. 1G). Additionally, the sequence homology within the ZIC ZF-NC suggests that all human ZICs may be SUMOylated at Motif 1.

### ZIC5 SUMOylation is critical in vivo to drive neural crest specification

To determine if SUMOylation alters ZIC5 function *in vivo*, denaturing HPLC was used to screen a genomic DNA library of ENU mutagenised BALB/c mice (Coghill *et al.*, 2002) for a mouse strain in which ZIC5 could not be SUMOylated. Thus, a genome was identified with an A to G transition in exon 1 of murine *Zic5* at position 1429 (NM_022987.3) that resulted in a K363R mutation within the ZIC5 ZF-NC (ZIC5 K363R) (Fig. 2A,B). The resulting allele was named kiska (*Ki*) and recovered from the frozen sperm archive. To uncover the effect of the mutation on ZIC5 function, the new allele was compared to an existing *Zic5* null line (*Zic5*^*−/−*^). Consistent with previous reports of *Zic5*^*−/−*^ mice (Inoue *et al.*, 2004), a proportion of *Zic5*^*−/+*^ animals developed hydrocephaly post-birth (5%, N=54; Fig. 2C). Additionally, it was observed that a proportion of *Zic5*^*−/+*^ animals (13%, N=123) exhibited a ventral spot (Fig. 2D), indicative of trunk NCC hypoplasia, which is consistent with previous reports that insufficient NCCs arise in *Zic5*^*−/−*^ embryos (Inoue *et al.*, 2004) and serves as a marker of trunk NCC depletion. *Zic5*^*Ki/+*^ mice also presented with a ventral spot (3%, N=73), although at a lower rate than the heterozygous nulls, suggesting that the mutation produces a hypomorphic allele. This was confirmed by placing the *Zic5 Ki* allele in *trans* to the null. Mice of the genotype *Zic5*^*Ki/-*^ exhibited increased penetrance of the ventral spot (29%, N=21, *p*<0.05 G-test) relative to the heterozygous null allele, as predicted for a hypomorphic allele. These results, in which SUMO-incompetent (i.e. unmodified) ZIC5 is sufficient to ameliorate but not fully rescue the null phenotype, indicate that the SUMOylated form of ZIC5 is critical during NCC development.

**Figure 2:**
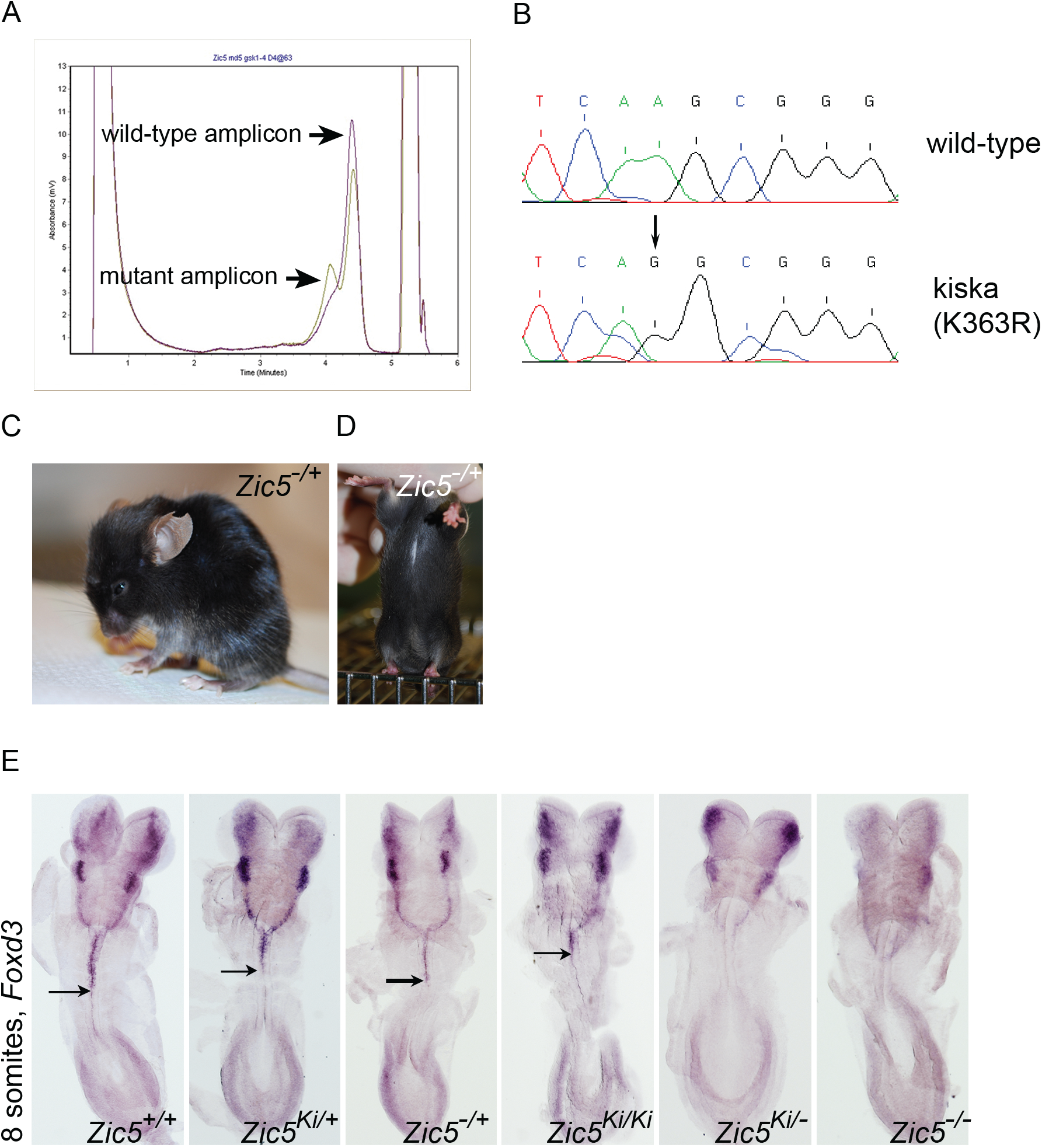
The K363R (kiska; Ki) hypomorphic allele of Zic5 leads to hypoplasia of trunk neural crest. (A) Denturing HPLC trace of the mutation-containing amplicon from pooled genomic DNA of four individual animals. (B) Sequence trace of wild-type and homozygous K363R animals, showing the A to G transition (arrow) at position 1429 of *Zic5* (of NM_022987.3). (C) Lateral view of *Zic5* mutant animal with hydrocephaly which was detected at low frequency in *Zic5*^*−/+*^ and *Zic5*^*Ki/−*^ animals. (D) Ventral view of *Zic5* mutant animal showing a ventral spot which was detected at low frequency in mice of the genotypes *Zic5*^*−/+*^, *Zic5*^*Ki/+*^ and *Zic5*^*Ki/−*^. (E) Dorsal view of intact 8 somite-stage embryos following WMISH to *Foxd3* (anterior to the top). Arrow indicates posterior limit of trunk neural crest *Foxd3* expression. Embryos of the genotypes *Zic5*^*Ki/+*^, *Zic5*^*−/+*^ and *Zic5*^*Ki/Ki*^ have reduced *Foxd3* expression and embryos of the genotypes *Zic5*^*Ki/−*^ and *Zic5*^*−/−*^ have ablated *Foxd3* expression compared to stage-matched wild-type (*Zic5*^*+/+*^) embryos. A minimum of four staged matched embryos per genotype were compared to precisely staged matched litter mates.

To better understand the role of ZIC5 SUMOylation in neural crest development, the expression of the neural crest specifier gene *Foxd3* was examined in embryos that lack ZIC5 (*Zic5*^*−/−*^) and those in which the SUMO-incompetent form of ZIC5 is present (*Zic5*^*Ki/Ki*^). Embryos expressing only the SUMO-incompetent form of ZIC5 have substantially depleted *Foxd3* expression (Fig. 2E), demonstrating that K363 modification is essential for the optimal transcription factor activity of ZIC5. The level of *Foxd3* expression is further depleted in embryos trans-heterozygous for the two alleles (*Zic5*^*Ki/−*^) or those that lack *Zic5* (*Zic5*^*−/−*^; Fig. 2E), indicating that the SUMO-incompetent form of ZIC5 retains some transactivation ability.

### SUMOylation promotes the transcriptional ability of ZIC5 at ZIC responsive elements

To understand how SUMOylation influences human ZIC5 function, cell based assays of ZIC transcription factor, co-factor and macro-molecular activity were employed. First, the consequence of ZIC5 SUMOylation for transactivation was assayed using an established *APOE*:Luc2 reporter assay (Ahmed *et al.*, 2020) which is based upon an initial observation by Salero *et al.* (2001) that ZIC1 and ZIC2 proteins are able to stimulate transcription via a genomic fragment from the human *APOE* promoter. The ZIC5 protein is also able to transactivate this promoter fragment in HEK293T cells, whereas a form of ZIC5 with a mutation within the DNA-binding domain V5-ZIC5-C528S is not (Ahmed *et al.*, 2020). These results are independently reproduced here. Additionally, a plasmid IP approach was used to verify that the V5-ZIC5-C528S protein does not interact with the *APOE* promoter fragment, thus confirming that ZIC5 transactivation of the *APOE* promoter in this assay is dependent upon DNA interaction (Fig. S2A, A’, B). When HEK293T cells were transiently transfected with the *APOE*:Luc2 reporter and the SUMO-incompetent V5-ZIC5-K393R, the transactivation of the promoter fragment was significantly reduced (but not ablated) relative to that stimulated by V5-ZIC5-WT (Fig. 3A, A’; *p*<0.05). Moreover, when SUMOylation of wild-type ZIC5 protein was prevented (by inhibition of universal SUMOylation via co-expression of a dominant negative form of UBC9; Flag-UBC9-C93S (Poukka *et al.*, 1999)) the transactivation ability of V5-ZIC5-WT was reduced to the level of V5-ZIC5-K393R (Fig. 3B, B’). The results indicate that SUMOylation of ZIC5 at K393 is necessary to drive maximal transactivation of the *APOE* promoter in HEK293T cells.

**Figure 3:**
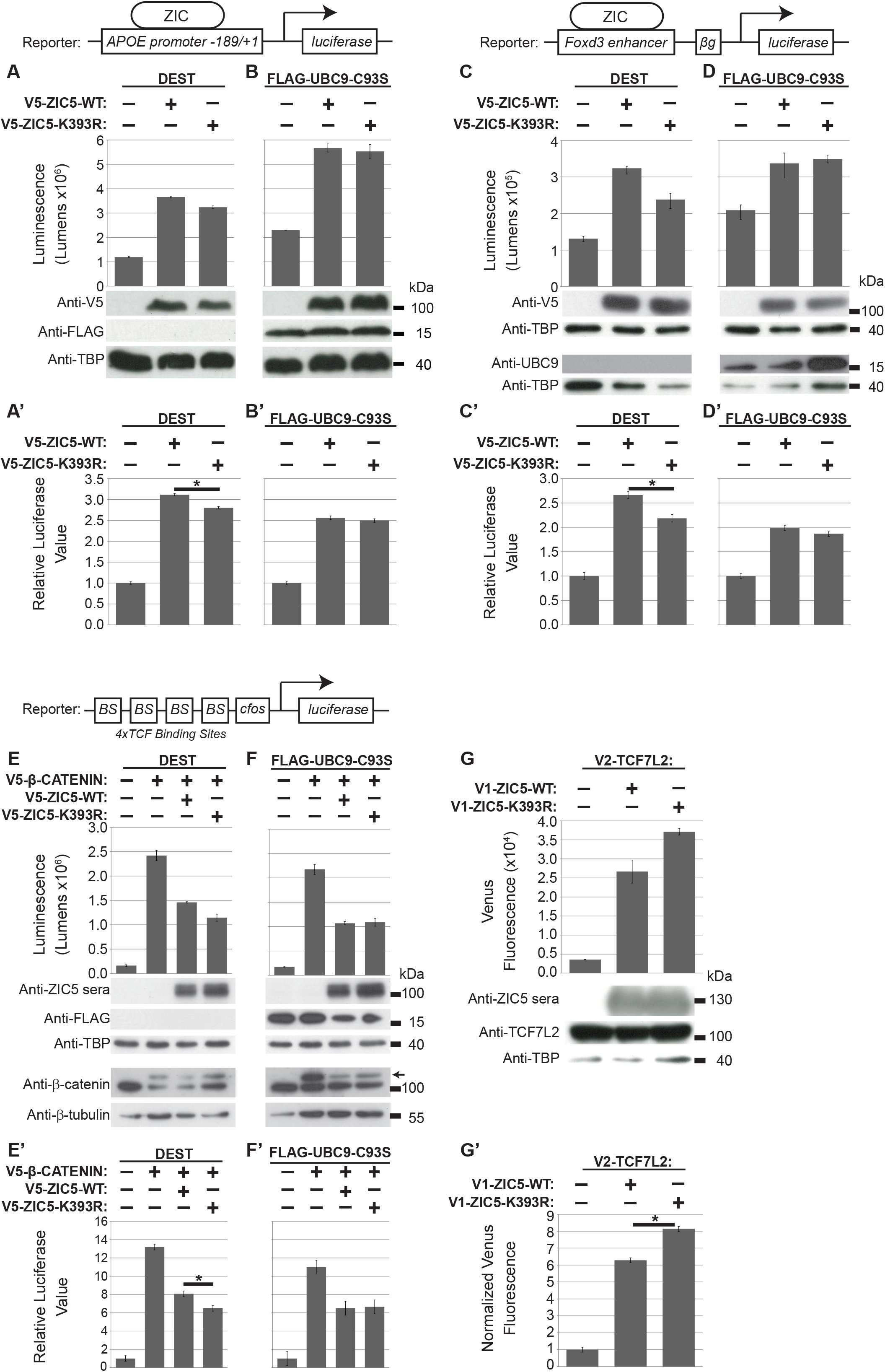
SUMOylation alters ZIC5 gene regulation activity and macro-molecular interaction. (A –F’) Luciferase reporter activity in HEK293T cells with the reporter and expression constructs indicated. (A and A’) The K393R SUMO-incompetent form of ZIC5 shows a significant decrease in transactivation ability compared to wild-type ZIC5. (B and B’) The transactivation ability of wild-type ZIC5 is impeded when SUMOylation is universally inhibited by UBC9-C39S and is equivalent to the SUMO-incompetent form of ZIC5. (C and C’) The K393R SUMO-incompetent form of ZIC5 shows a significant decrease in transactivation ability. (D and D’) The transactivation ability of wild-type ZIC5 is impeded by universal SUMOylation inhibition via UBC9-C39S and is equivalent to that of the SUMO-incompetent form of ZIC5. (E and E’) The K393R SUMO-incompetent form of ZIC5 shows a significant increase in inhibition of β-catenin mediated transcription relative to that obtained with wild-type ZIC5. (F and F’) The ability of wild-type ZIC5 to inhibit β-catenin mediated transcription is increased to a level equivalent to that of the K393R SUMO-incompetent form. (G and G’) BiFC assay in HEK293T cells. The K393R SUMO-incompetent form of ZIC5 shows increased interaction with TCF7L2 relative to the wild-type form of ZIC5. (A-G) Raw data and WB to show overexpressed proteins from one representative experiment. Error bars = SD from three internal repeats. (A’-G’) Pooled data from three external repeats (normalised to the background). Error bars = SEMs (ANOVA). *: *p*<0.05, two-way ANOVA with Bonferroni multiple comparison test.

To determine whether the same mechanism may contribute to the decreased *Foxd3* expression observed in the *Ki* SUMO-incompetent mouse strain (Fig. 2E), this set of experiments were repeated using a recently generated *Foxd3*:Luc2 reporter assay (Ahmed *et al.*, 2020). This construct incorporates the mouse genomic region equivalent to the recently identified ZIC responsive chick *Foxd3* enhancer (Simões-Costa *et al.*, 2012). The wild-type ZIC5 protein is able to transactivate this promoter fragment in HEK293T cells, whereas a form of ZIC5 with a mutation within the DNA-binding domain V5-ZIC5-C528S is not (Ahmed *et al.*, 2020). These results are independently reproduced here. Additionally, a plasmid IP approach was used to verify that the V5-ZIC5-C528S protein does not interact with the *Foxd3* enhancer fragment, thus confirming that ZIC5 transactivation of the *Foxd3* element in this assay is dependent upon DNA interaction (Fig. S2C, C’, D). Luciferase assays with the *Foxd3*:Luc2 reporter in which SUMOylation of ZIC5-K393 was inhibited specifically (via K393R substitution) or universally (via co-expression of Flag-UBC9-C93S) recapitulated the results obtained with the *APOE:*Luc2 reporter (Fig. 3C, C’, D, D’; *p*<0.05). The data indicate that SUMOylation at K393 is necessary to maximise transactivation of the *Foxd3* enhancer element. To evaluate the role of TCF7L2 in *Foxd3* expression, TCF7L2 was expressed alone or in conjunction with wild-type ZIC5. In contrast, the co-expression of FLAG-TCF7L2 with V5-ZIC5-WT did not increase transactivation of the *Foxd3*:Luc2 reporter (Fig. S2E), suggesting TCF does not co-operate with mammalian ZIC protein at ZREs and indicates a specific requirement for ZIC5 SUMOylation.

### SUMOylation decreases ZIC5 co-inhibition of WNT signalling

To examine the effect of ZIC5 SUMOylation on co-factor activity, the ability of ZIC5 and ZIC5-K393R to inhibit WNT/β-catenin dependent transcription was examined by a TOPflash reporter assay that specifically measures β-catenin/TCF mediated transcription (Korinek *et al.*, 1997). As shown in Fig. S3A, A’ when this assay is conducted in HEK293T cells using our standard protocols, transfection of V5-β-CATENIN stimulates expression from the TOPflash but not the control FOPflash reporter, and co-transfection with the V5-ZIC5-WT expression vector inhibits expression from the TOPflash (but does not alter expression from the FOPflash) reporter. To evaluate the influence of SUMOylation on ZIC5-mediated WNT inhibition, SUMOylation was inhibited either specifically (via expression of the SUMO-incompetent V5-ZIC5-K393R construct; Fig. 3E, E’) or globally (via co-expression of Flag-UBC9-C93S with V5-ZIC5-WT; Fig. 3F, F’). Overexpression of V5-ZIC5-K393R significantly enhanced suppression of β-catenin stimulated transcription relative to V5-ZIC5-WT (*p*<0.05) and the presence of dominant negative UBC9 converted V5-ZIC5-WT to a more efficient inhibitor of β-catenin mediated transcription. Together these assays indicate that the unmodified form of ZIC5 is required for optimal co-factor activity and that SUMOylation decreases the co-factor activity of ZIC5.

### SUMOylation reduces formation of the ZIC5/TCF repression complex

SUMOylation is known to alter macro-molecular interactions (Geiss-Friedlander and Melchior, 2007; Flotho and Melchior, 2013; Matunis and Rodriguez, 2016). The possibility that K393 SUMOylation affects the previously characterised ZIC/TCF interaction (Pourebrahim *et al.*, 2011; Fujimi, Hatayama and Aruga, 2012) was considered a potential cause of the SUMOylation induced change in ZIC5 behaviour (i.e. from inhibition at WREs towards transactivation at ZREs). The physical interaction between TCF7L2/ZIC5-WT and between TCF7L2/ZIC5-K393R was examined via bimolecular fluorescence complementation (BiFC) assay (Kodama and Hu, 2012) using a split Venus fluorescent molecule (V1 and V2). As shown in Fig. 3G, the SUMO-incompetent form of ZIC5 (V1-ZIC5-K393R) exhibited increased interaction with TCF7L2 (V2-TCF7L2) relative to ZIC5-WT (V1-ZIC5-WT). This interaction was validated though a BiFC competition assay where wild-type ZIC5 protein without the V1 tag competes with V1-ZIC5, indicating a specific interaction (Fig. S3B). This is consistent with both the improved ability of V5-ZIC5-K393R to inhibit β-catenin/TCF mediated transcription and the decreased ability of V5-ZIC5-K393R to drive expression at ZREs. In contrast, our experiments found no support for the alternative possibility that SUMOylation altered ZIC5 subcellular localisation (Fig. S4A-C). As lysine residues can be the target of PTMs other than SUMOylation, the ability of K393 to undergo ubiquitylation was assessed using an Ubiquitin-based BiFC assay to further explore the possibility that another K393 PTM is responsible for the observed effects on ZIC5 transcriptional control. Although ZIC5 was found to be ubiquintated, K393 is not a target lysine (Fig. S4D).

### The proportion of SUMOylated ZIC5 protein varies with the strength of canonical WNT signal

If, as indicated by the TOPflash and BiFC assays, SUMOylation drives the demise of the TCF/ZIC repressor complex, then it may be expected that the proportion of ZIC5 that is SUMOylated varies with the strength of WNT signal. To test this possibility, the proportion of SUMOylated ZIC5 was compared in HEK293T cells in their basal state and in the presence of the GSK3 (a core component of the WNT signalling network) inhibitor LiCl. As shown in Figs. 4A, A’ and S5A, activation of WNT signalling via LiCl caused an increase in the proportion of SUMOylated ZIC5. A TOPflash assay confirmed that the LiCl treatment increased WNT/β-catenin-mediated transcription as expected (Fig. S5B). Additionally, hyper-stimulation of WNT signalling via LiCl decreased the ability of V5-ZIC-WT to inhibit WNT/β-catenin-mediated transcription in a time dependent manner (Fig. 4B). This indicates that as WNT signalling activity increases (and the proportion of SUMOylated ZIC5 increases) the ability of ZIC5 to inhibit TCF-dependent transcription decreases. Together these experiments confirm WNT signalling-dependent ZIC5 SUMOylation results in a shift in ZIC5’s transcription regulation function from inhibitory co-factor at WREs to transcriptional activator at ZREs.

**Figure 4:**
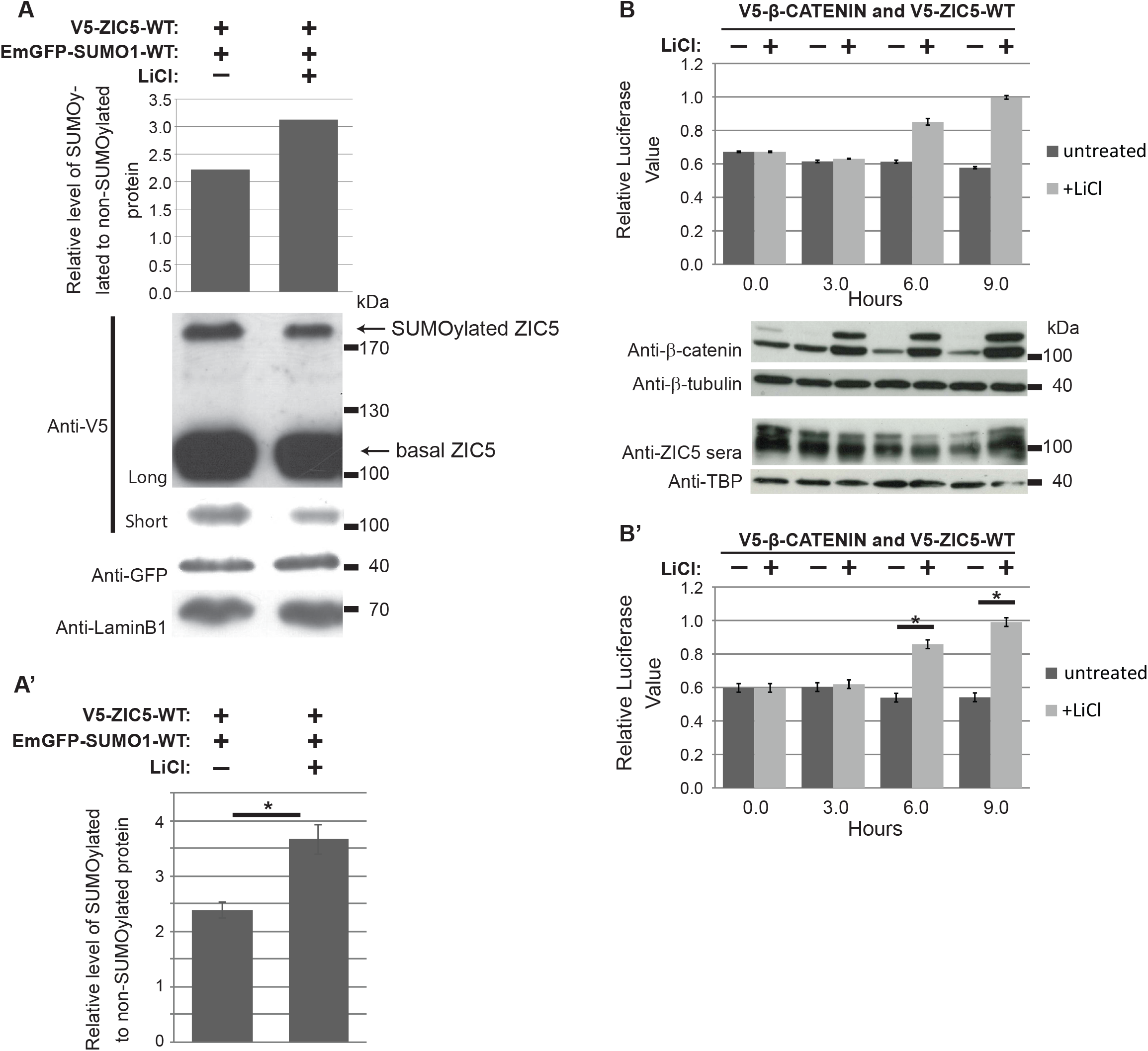
Activation of the canonical WNT pathway promotes ZIC5 SUMOylation and TCF-dependent transcription. (A and A’) Relative quantification of the proportion of SUMOylated ZIC5 based on WB analysis of ZIC5 protein from HEK293T cells co-transfected with V5-ZIC5-WT and EmGFP-SUMO1-WT, followed by incubation in the presence or absence of LiCl. Basal ZIC5 was quantified from the short exposure and SUMOylated ZIC5 from the long exposure. (A) Representative WB and corresponding quantification. (A’) Average relative levels from four external repeats. Error bars = SEM, *: *p*<0.01 paired *t*-test. (B and B’) Luciferase reporter activity in HEK293T cells using the TOPflash reporter construct in the presence or absence of LiCl over a 9 hr time-course. Note that the untreated normalised Relative Luciferase Value is less than 1, indicative of V5-ZIC5-WT acting as an inhibitor of β-catenin dependent transcription. LiCl treatment causes this value to rise, consistent with a loss of ZIC5 inhibitory activity (B) Representative experiment with corresponding WB of overexpressed proteins. Error bars = SD of three internal repeats. (B’) Relative luciferase values from three external repeats. Error bars = SEM of three external repeats. *: *p*<0.05, two-way ANOVA with Bonferroni multiple comparison test. In both cases data is normalised to the V5-DEST/β-catenin transfection corresponding to each sample.

## Discussion

This study shows that the multifunctional transcription regulator, ZIC5, is a SUMO substrate, being poly-SUMOylated at a conserved lysine residue within the ZF-NC domain (K393). The SUMOylation state of ZIC5 shifts the balance of ZIC5 function: increased SUMOylation correlates with a decreased propensity to interact with TCF and repress WREs and an increase in transactivation at ZREs. Additionally, elevated canonical WNT signalling is associated with a higher proportion of SUMOylated ZIC5 protein and a decrease in the ability of ZIC5 to inhibit transcription at WREs. The results of the cell-based assays are synthesized in a working model (Fig. 5) that illustrates how a high canonical WNT environment and demise of the ZIC/TCF repressor complex can facilitate transactivation at both WREs and ZREs. The *Ki* mouse strain, in which the conserved lysine within the ZF-NC cannot be SUMOylated, exhibits the same phenotype as the complete loss of the ZIC5 protein, albeit in a less severe/frequent manner. When placed in *trans* to the null allele, the phenotype is further enhanced, indicating that *Ki* is a partial loss-of-function allele. This demonstrates that basal ZIC5 is insufficient to fully perform the activities of the ZIC5 protein and that PTM of this conserved lysine is required *in vivo*. The work also provides a direct demonstration that PTM of this residue is necessary to drive optimal *Foxd3* expression and NCC specification.

**Figure 5:**
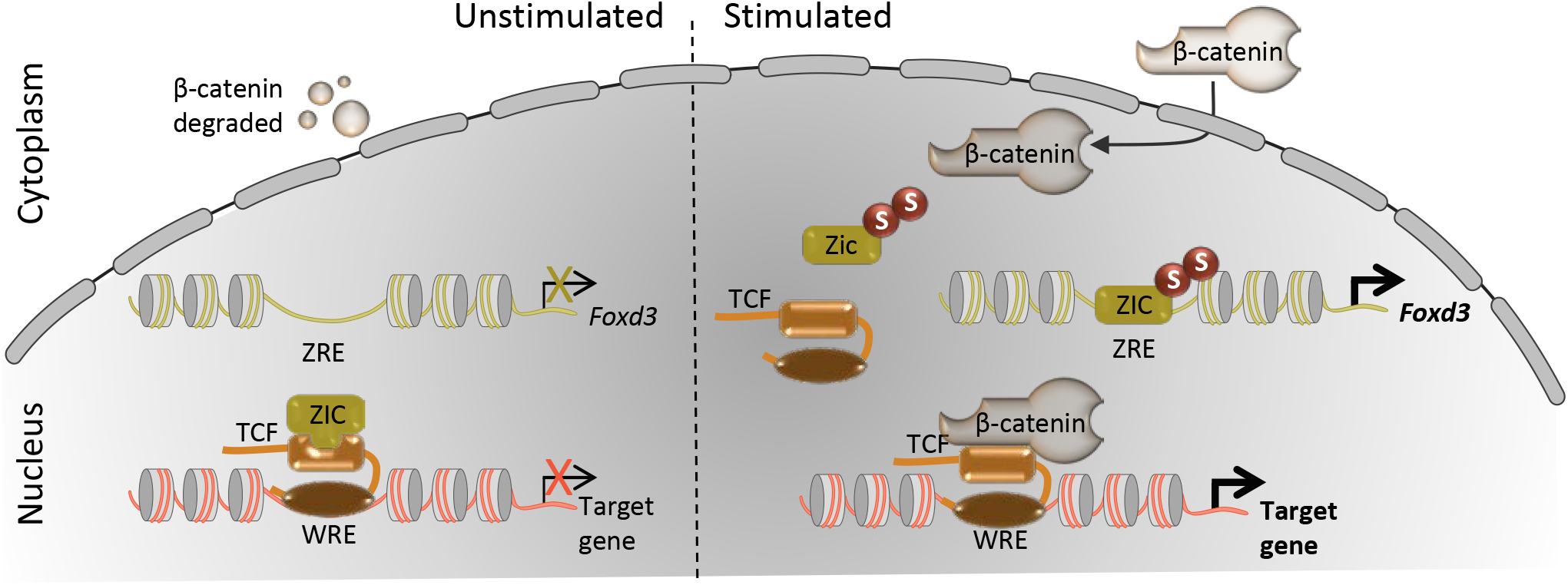
WNT responsive SUMOylation of ZIC5 influences both TCF */β*-catenin dependent and independent gene expression. Prior to signalling (left: unstimulated state), WNT target genes can be constitutively inhibited by nuclear TCF which recruits transcriptional co-repressors such as ZIC proteins to WNT responsive elements (WRE). This recruitment could limit the availability of ZIC protein and prevent activation at ZIC responsive elements (ZRE). In this state, cytoplasmic β-catenin is degraded by the cytoplasmic destruction complex. WNT ligand binding (right: stimulated state) initiates a cascade of cytoplasmic events (not shown) culminating in β-catenin nuclear entry and also drives SUMOylation (S) of ZIC to promote the dissociation of the ZIC/TCF repressor complex. This facilitates both the activation of gene expression via ZREs and the formation of the TCF/β-catenin complex to activate expression at WREs.

SUMOylation is a dynamic and reversible PTM that occurs at a lysine residue found within a consensus protein sequence (Han *et al.*, 2018). All mammalian ZIC proteins contain at least one such consensus sequence immediately N-terminal of the zinc finger domain. The deep conservation of this ZIC protein region has been highlighted in multiple phylogenetic ZIC analyses (Aruga *et al.*, 2006; Tohmonda *et al.*, 2018), but its functional significance remains unknown. Here we show that ZIC5 can be SUMOylated at K393 in HEK293T cells and that conservative substitution of this lysine residue with the SUMO-incompetent arginine residue is sufficient to prevent all ZIC5 SUMOylation (Fig. 1D). In contrast, individual mutation of the two other ZIC5 high probability SUMO target lysines does not alter the pattern of SUMO-dependent protein products (Fig. 1C). We conclude that K393 is the sole *in vivo* target of SUMO and furthermore that, based on the size of the dominant SUMO-dependent products, ZIC5 is poly-SUMOylated at K393. Our conclusion is supported by high throughput proteomic studies which have identified ZIC5 as a SUMO substrate (but which did not identify the target lysine residue) (Hendriks and Vertegaal, 2016). Our finding also extends the work of Chen *et al.*, (2013) who demonstrated that the paralogous lysine within human ZIC3 (K248) can be SUMOylated in HeLa cells.

When overexpressed in HeLa cells, ZIC3 SUMOylation was found to influence nuclear localisation. Generally, studies of ZIC sub-cellular distribution report predominate nuclear localisation (Koyabu *et al.*, 2001; Ishiguro *et al.*, 2004; Ware *et al.*, 2004; Brown *et al.*, 2005; Ahmed *et al.*, 2013, 2020), and preventing ZIC3 SUMOylation leads to diffuse subcellular distribution. In contrast, preventing ZIC5 SUMOylation did not alter its subcellular distribution (Fig. S4A-C). Instead, we found that SUMOylation decreased interaction with TCF7L2 and ZIC inhibition at WREs as well as increasing trans-activation at ZREs. The findings further suggest that the common consequence of SUMOylation is to alter protein-protein interactions. Additionally, we observed that the kinetics of ZIC5 SUMOylation vary with the level of WNT activity (Fig. 4). We speculate that the need to switch between different modes of ZIC transcription control in response to inductive signals could explain the evolutionary conservation of the ZF-NC domain. In support of this idea, it is noteworthy that the *C. elegans* single ZIC orthologue REF-2 lacks both a recognizable ZF-NC domain and consensus SUMOylation site, and exhibits a distinct mode of operation in which TCF acts as a co-factor to promote expression at ZREs (Murgan *et al.*, 2015). If the ZF-NC conservation is driven by SUMOylation at the conserved lysine within this domain, we would predict that the other mammalian ZIC proteins are also SUMOylated at the paralagous and orthologous lysine.

ZIC proteins are known to inhibit WNT signalling when overexpressed in cells, and in *Xenopus* and zebrafish embryos (Pourebrahim *et al.*, 2011; Fujimi, Hatayama and Aruga, 2012; Zhao *et al.*, 2019). Here we show that ZIC5 also inhibits β-catenin/TCF-dependent transcription, measured via TOPflash assay in HEK293T cells. Strikingly, hyperstimulation of canonical WNT signalling in this assay, via LiCl, led to a loss of ZIC5 inhibition (Fig. 4B). This implies ZIC inhibition of transcription at WREs is context dependent: robust in a relatively low WNT environment, but overcome in the presence of sustained WNT signalling. Given that sustained WNT signalling promotes ZIC5 SUMOylation and that ZIC5 SUMOylation decreases the ZIC5/TCF7L2 interaction and repression at WREs, it is plausible that SUMOylation contributes to the observed context dependent ZIC5 transcriptional regulation. This is supported by the fact that TCF7L2 (formerly known as TCF4) can be SUMOylated, and modification enhances the β-catenin-dependent transcriptional activation of TCF7L2 (Yamamoto *et al.*, 2003). It identifies another SUMOylated WNT pathway component, further strengthening the observation that SUMOylation is a core regulator of canonical WNT signalling (Kim, Chia and Costantini, 2008; Choi *et al.*, 2011; Gao, Xiao and Hu, 2014). As shown in Fig. 5, this feature of ZIC/TCF co-repression enables sustained WNT signalling to direct trans-activation not just at WREs but also at ZREs. The *in vivo* relevance of this model is demonstrated by the finding that prevention of ZIC5 SUMOylation is sufficient to decrease *Foxd3* expression, previously shown to require ZIC binding to an upstream enhancer, during murine NCC specification. Furthermore, cis-regulatory analysis has failed to identify many WREs critical for expression of neural crest specifiers (Simões-Costa and Bronner, 2015) and the work presented here demonstrates how canonical WNT signalling can drive NCC specification in the absence of such WREs.

One caveat of the work presented here is that the role of canonical WNT signalling in mouse NCC specification remains ambiguous. For example, conditional deletion of many candidate WNT genes does not prevent murine NCC development. It is however possible that this is due to technical reasons relating to the conditional gene deletion strategy most often used in the mouse (Barriga *et al.*, 2015). Indeed, the use of an alternative *Cre* driver has provided evidence that ectopic WNT signalling drives NCC development in the murine forebrain (Mašek *et al.*, 2016), demonstrating that WNT signalling is able to induce murine NCC specification. Our work provides insight into how the putative WNT signals can restrict the NCC precursor domain within the NPB, despite a broader domain of expression of NCC specifying transcription factors such as the ZIC genes. High WNT activity at the NPB (Ferrer-Vaquer *et al.*, 2010) will simultaneously convert both ZIC and TCF proteins into transactivators, meaning that target NCC specifier genes under the control of either a ZRE (such as demonstrated for the *Foxd3* enhancer; Fig. 3C, C’ and Fig. S2E, E’) or WRE can be activated. Simultaneously, in the future lateral neurectoderm (where WNT signals are lower (Ferrer-Vaquer *et al.*, 2010) and ZICs and TCFs are co-expressed; K.S.B and R.M.A unpubished data), TCF/ZIC co-repression at WREs and low occupancy at ZREs will persist.

The work here focused on SUMOylation of ZIC5, driven by the knowledge that SUMOylation appears important during chick and *Xenopus* NCC development. In chick, the SUMO ligase is expressed specifically in the neural crest and SUMOylation of the NPB transcription factor *Pax7* is required for full expression of the specifying transcription factors *Snail2*, *Sox9* and *Foxd3* (Luan *et al.*, 2013). Additionally, some specifying transcription factors (Sox9 and Sox10) are themselves SUMOylated in chick and *Xenopus* embryos and it will be of interest to determine whether WNT driven SUMOylation at the NPB provides a concerted method of directing NCC specification. We are the first to implicate SUMOylation in mouse neural crest development: we demonstrates a clear requirement for ZIC5 K393 to elicit full expression of *Foxd3* during NCC formation. Additionally, the cell-based assays demonstrate that SUMOylation of this residue is also required for full expression of *Foxd3* in HEK293T cells. Given that several studies have described the overlap of SUMO sites with other PTMs (such as ubiquitylation, acetylation and methylation), it remains possible that some other ZIC5 K393 PTM contributes to the observed effect in the *Ki* mouse. The most common overlap occurs between ubiquitylation and SUMOylation (Hendriks and Vertegaal, 2016) and, despite ZIC5 being ubiquitylated, K393 is not an ubiquitin target (Fig. S4D). Understanding whether other PTM of K393 occurs awaits further characterisation of ZIC5 PTMs.

Overall, the data presented here are consistent with a mechanism by which, in a high canonical WNT environment, PTM of the ZIC5 protein alters the balance between alternative modes of Zic transcription control *in vitro* and *in vivo*. This mechanism appears important during WNT-induced NCC specification. Furthermore, the proposed consequence of this, whereby WNT signals alter transcription at elements other than canonical WREs, is potentially of broad significance and may be used in many other signalling events and pathways. The use of hetero-protein complexes to repress transcription enables a nuclear store of synthesised, but inactive, transcription factors ready to regulate transcription in response to dynamic extracellular cues.

## Experimental procedures

### Expression and reporter construct generation

The generation of pCMV6-XL5-ZIC5, pENTR3C-ZIC5-WT, V5-ZIC5-WT, TOPflash and FOPflash (containing a *cfos* promoter and four wild-type or mutant TCF binding sites) has been previously described (Ahmed *et al.*, 2013). Overlap extension PCR was used to introduce the K393R mutation into pCMV6-XL5-ZIC5 and a FseI/BstEII fragment of human *ZIC5* digested from the mutated pCMV6-XL5-ZIC5 was used to replace the equivalent region in pENTR3C-ZIC5-WT to create pENTR3C-ZIC5-K393R. The K522R mutation was introduced into pENTR3C-ZIC5-WT by recombineering to create pENTR3C-ZIC5-K522R. Overlap extension PCR was used to introduce the C528S mutation within pENTR3C-ZIC5-WT to generate pENTR3C-ZIC5-C528S. To generate UBC9-fused proteins for the UFDS assay, the UBC9 cDNA (with stop codon deleted) was amplified from pSG5-HA-hUBC9 (Chang *et al.*, 2007) and inserted into the KpnI restriction enzyme site at the *ZIC5* N-terminus to create pENTR3C-UBC9-ZIC5-WT, pENTR3C-UBC9-ZIC5-K393R and pENTR3C-UBC9-ZIC5-K522R. In each case, the insert from the entry clone was transferred to the destination clone pcDNA3.1/nV5-DEST (Life Technologies) or V1-ORF-DEST (see below) via a Gateway LR Clonase reaction (as per manufacturer’s instructions; Life Technologies) to produce the following plasmids: V5-ZIC5-K393R, V5-ZIC5-K522R, V5-UBC9-ZIC5-WT, V5-UBC9-ZIC5-K393R, V5-UBC9-ZIC5-K522R, V1-ZIC5-WT and V1-ZIC5-K393R.

To generate pENTR3C-SUMO1-WT, human *SUMO1* cDNA was PCR amplified from pEYFPC3-SUMO-1 (Harder, Zunino and McBride, 2004) and cloned into BamHI/XhoI restricted pENTR3C. The SUMO-defective mutant (pENTR3C-SUMO1-ΔGG) was designed based on information in (Kamitani, Nguyen and Yeh, 1997), and the cDNA synthesised and cloned into pENTR3C by GeneScript. In each case, the insert from the entry clone was transferred to the destination clone Vivid Colors pcDNA 6.2/N-EmGFP-DEST (Life Technologies) via a Gateway LR Clonase reaction to produce the following plasmids: EmGFP-SUMO1-WT and EmGFP-SUMO1-ΔGG.

To generate pENTR3C-TCF7L2, human *TCF7L2* (previously called *TCF4*) cDNA was PCR amplified from pFLAG-TCF4 (Pourebrahim *et al.*, 2011) and cloned into EcoRI/KpnI restricted pENTR3C. The insert was transferred to the destination vector V2-ORF-DEST (see below) via a Gateway LR Clonase reaction to create V2-TCF7L2.

The V1-ORF-DEST and V2-ORF-DEST expression constructs contain the N-terminal CDS of the Venus protein (designated V1) or C-terminal CDS of the Venus fluorescent protein (designated V2) upstream of a site suitable for destination cloning. When used in a Gateway LR Clonase reaction with an entry construct (such as pENTR3C), a mammalian expression construct is generated that expresses a fusion protein (either V1- or V2-) and the protein encoded within the entry construct. For example, V2-Ubiquitin contains the V2 C-terminal fragment fused to the CDS of Ubiquitin.

The other expression constructs have been previously described: V5-β-CATENIN (pcDNA3.1/V5-HisA-β-CATENIN) (Usami *et al.*, 2003) and pFLAG-UBC9-C93S (Poukka *et al.*, 1999).

The *APOE* and *Foxd3* reporters have been described previously as B:*luc2*:APOE and B:*luc2*:Foxd3, respectively (Ahmed *et al.*, 2020).

### Cell culture

The Human Embryonic Kidney cell line (HEK293T) was cultured and transiently transfected as previously described (Ahmed *et al.*, 2013).

### Subcellular fractionation, SDS-PAGE and Western Blotting (WB)

Subcellular fractionations were obtained via a nuclear protein extraction kit (Pierce NE-PER kit) according to the manufacturer’s protocol with the following modifications: 4 × 100 mm tissue culture (TC) dishes (approximately 2.8 × 10^7^ cells, for non-UFDS experiments, Corning®; cat. no. CLS430167), 1 × 60 mm TC dish (approximately 2.5 × 10^6^ cells, for UFDS experiments, Corning®; cat. no. CLS430166) or 6-well TC plates (approximately 1 × 10^6^ cells, for luciferase reporter assay WB, Costar®, cat. no. CLS3516) of 90-100% confluent HEK293T cells were lysed directly using CERI and NER buffer supplemented with 1x protease inhibitor cocktail (Roche), 1x PhosSTOP (Roche), 2 mM iodoactemide, and 1.6 mM N-Ethylmaleimide. 2 mM dithiothreitol (DTT) (Sigma Aldrich) and 1x NuPAGE LDS Sample Buffer (Life Technologies) were added to nuclear and cytoplasmic fractions and then heated for 5 min at 90°C. Samples were then loaded onto 7% or 12% SDS-PAGE gels and run at 100 V until proteins were sufficiently separated. Proteins were transferred to PVDF membranes (Millipore) via wet transfer at 15 V for 16 hrs. Membranes were blocked overnight at 4°C with 5% skim milk powder/0.2% Tween 20 (Sigma Aldrich)/PBS solution (WB blocking buffer) before being immunoblotted using standard WB techniques. To detect protein bands, blots were incubated with SuperSignal West Pico reagent (Pierce) then exposed to film (Amersham Hyperfilm MP, GE Life Sciences). Developed films were scanned and assembled in Adobe Illustrator CS5.1.

### Antibodies

The following primary antibodies were used: mouse monoclonal anti-V5 (1:200 dilution IF, 1:5000 dilution WB, Life Technologies, cat. no. R960-25), rabbit polyclonal anti-GFP (1:1000 dilution IF, 1:1500 dilution WB, Cell Signaling, cat. no. 2555), rabbit polyclonal anti-lamin B1 (1:1000 dilution IF, 1:1500 dilution WB, Abcam, cat. no. ab16048), mouse monoclonal anti-β-tubulin (1:1000 dilution WB, Abcam, cat. no. ab7792), mouse monoclonal anti-TATA binding protein (TBP; 1:2000 dilution WB, Abcam cat. no. ab818), goat polyclonal anti-β-catenin C-18 (1:500 dilution WB, Santa Cruz, cat. no. sc-1496), mouse monoclonal anti-UBC9 (1:1000 dilution WB, BD, cat. no. 610748), mouse monoclonal anti-TCF7L2 (1:1000 dilution WB, Abcam, cat. no. ab32873), mouse monoclonal anti-FLAG (1:1000 dilution WB, Sigma, cat. no. F1804) and rabbit anti-ZIC5 sera (1:500 dilution WB). The ZIC5 antibody was generated in rabbits using the epitope described in Inoue *et al*, (2004) as an antigen using standard techniques. The following secondary antibodies were used for immunofluorescence (1:500 dilution): Alexa^594^ and Alexa^488^ conjugated donkey anti-mouse (Molecular Probes, cat. no. A21206) and anti-rabbit (Molecular Probes, cat. no. A21203). The following secondary antibodies were used for WB (1:5000 dilution): horse radish peroxidase (HRP)-conjugated rabbit anti-mouse, rabbit anti-goat, and goat anti-rabbit (Zymed, Life Technologies). All antibodies were diluted in blocking buffer.

### Plasmid Immunoprecipitation (pIP) and quantitative PCR

Plasmid Immunoprecipitation was performed as previously described (Ahmed *et al.*, 2020). HEK293T cells, grown in 100 mm TC dishes (Sigma; CLS430167) were transfected with 8 μg of *APOE* or *Foxd3* reporter construct and 16 μg of V5-ZIC5-WT or V5-ZIC5-C528S.

Quantitative PCR was performed as described in Ahmed *et al.*, (2020). The primers used for the *APOE* promoter were Ark1669 (5’-GACTGTGGGGGGTGGTC −3’) and Ark1670 (5’-AGACTTGTCCAATTATAGGGCTC −3’). Primers used to amplify the *Foxd3* region were Ark1671 (5’-GTACATTCAAGCTCCGTTGCC −3’) and Ark1672 (5’-CCAGAACCAGGCTCTAAATTGG −3’).

### Luciferase reporter assays

HEK293T cells, grown in 6-well TC plates (Costar; CLS3516), were transfected with the combination of constructs appropriate for each experiment. For ZIC transactivation assays a total of 4.5 μg of DNA was added per well: 1 μg of the *APOE* or *Foxd3* reporter construct, 3 μg of either the ZIC5 expression construct or the empty construct (pcDNA3.1/nV5-DEST™) and 0.5 μg of either empty pcDNA3.1/nV5-DEST vector or FLAG-DN-UBC9. For β-catenin-mediated transcription assays, a total of 4.5 μg of DNA was transfected per well: 1 μg of the TOPflash or FOPflash reporter vectors, either 1 μg V5-β-CATENIN expression construct, 2 μg of the appropriate ZIC5 expression construct or the empty pcDNA3.1/nV5-DEST vector, and 0.5 μg of either empty pcDNA3.1/nV5-DEST vector or FLAG-DN-UBC9. To assess WNT background levels, the 1 μg V5-β-CATENIN expression construct was substituted with 1 μg pcDNA3.1/nV5-DEST. 5.5-8 hr post-transfection, cells were dissociated from the growth surface using 0.05 g/L trypsin and plated in triplicate onto a solid white, TC treated, 96-well plate (Costar; CLS3917). To avoid positional bias of the luminometer, sample order was randomised for each independent experimental repeat. The remaining cells were re-plated for WB analysis. 24 hr post-transfection, the cells for the luciferase assay were exposed to 100 μL of a 1:1 dilution of DMEM and luciferase substrate (ONE-Glo Luciferase Assay System, Promega), and luminescence from each well measured in a GloMax-96 Microplate Luminometer (Promega). The cells reserved for WB were lysed.

### BiFC assays

HEK293T cells, grown in 6-well TC plates (Costar; CLS3513) were transfected with 2 μg of V2-TCF7L2 or V2-Ubiquitin and 2 μg of either V1-ORF-DEST, V1-ZIC5-WT or V1-ZIC5-K393R. For the competition assays, cells were transfected with 1 μg V2-TCF7L2, 1 μg of V1-ORF-DEST or V1-ZIC5-WT, and either 0 μg, 1 μg or 2 μg of V5-ZIC5-WT as well as pcDNA3.1/nV5-DEST to keep the total amount of DNA consistent. 24 hr post-transfection, cells were dissociated from the growth surface using 0.05 g/L trypsin and plated in triplicate onto a solid white, TC treated, 96-well plate (Costar; CLS3917). To avoid positional bias of the luminometer, sample order was randomised for each independent experimental repeat and the remaining cells re-plated for WB analysis. 48 hr post-transfection, media was removed from the cells for BiFC analysis, replaced with 1x PBS, and fluorescence measured using the M1000 Pro multimode fluorescence plate reader (Tecan). The cells reserved for WB were lysed.

### LiCl treatment

For WB analysis of the SUMOylated form of ZIC5, 4 × 100 mm tissue TC dishes (Corning; CLS430167), each containing approximately 5.6 × 10^7^ cells, were transfected with 12 μg of V5-ZIC5-WT and 12 μg of EmGFP-SUMO1-WT per plate. 6 hr later, cells were dissociated from the growth surface using 0.05 g/L trypsin, and replated into 8 × 100 mm TC dishes. 24 hr post-transfection, LiCl (final concentration 20 mM) was added to half of the dishes. 48 hr post-transfection, cells were lysed for WB. Post-WB, relative amounts of SUMOylated and non-SUMOylated protein were quantified from scanned images using ImageJ (NIH).

For TOPflash assays, HEK293T cells, grown in 100 mm tissue TC dishes (Corning; CLS430167), were transfected with TOPflash (6 μg), V5-β-CATENIN (6 μg) and either V5-DEST or V5-ZIC5-WT (12 μg). 6 hr post-transfection, cells were replated into 12-well dishes (Corning; CLS3513). 20-22 hr post-transfections, half of the dishes were treated with LiCl (final concentration 20 mM) before harvesting at 0, 1.5, 3, 6 and 9 h. At each time point ~6.0 × 10^4^ cells (from each treatment) were used to assay reporter activity and the remainder lysed for WB analysis.

### Immunofluorescence staining, microscopy and quantitation of subcellular localisation

Cells were prepared for immunofluorescence microscopy and the cytoplasmic and nuclear localisation was analysed as described previously (Ahmed *et al.*, 2013). For co-localisation experiments, cells were viewed using a Leica TCA SP5 confocal laser scanning microscope using a 63x oil immersion objective. The ImageJ (NIH) Line Tool was used to determine whether ZIC5 was enriched in SUMO1 foci. At least 120 SUMO1 foci were analysed for each experiment and three independent experiments were performed. Images for publication were assembled in Photoshop CS7.

### Isolation of the Zic5R (kiska; Ki) strain

The following primers were used to screen an archive of genomic DNA constructed from the F1 progeny of BALB/c N-ethyl-N-nitrosourea (ENU) mutagenised males and C3H/HeH females (Coghill *et al.*, 2002; Quwailid *et al.*, 2004): Ark280 (exon 1; Forward, 5’ CTT TCC TGC GCT ACA TGC 3’) and Ark281 (intron 1/2; Reverse, 5’ CAG GGA AAA ATG AAA GCG AAC 3’). The 435 bp fragment, spanning the ZF-NC domain and zinc fingers 1-3, was amplified from 5760 animals (arranged as 1440 pools, with each pool containing genomic DNA from four individual animals). DNA pooling, PCR and heteroduplex formation was as previously described (Quwailid *et al.*, 2004). Each amplicon was analysed by denaturing high performance liquid chromatography (DHPLC) on a Transgenomic Wave machine according to the manufacturer’s instructions (Transgenomic). For amplicons exhibiting a DHPLC trace divergent to that obtained from wild-type F1 DNA, the corresponding four DNAs were individually amplified, purified and subjected to Sanger Sequencing using standard procedures to discern the nature of the mutation and the identity of the carrier animal. Subsequently, the *Zic5 Ki* strain was recovered by IVF of C3H/HeH eggs, using standard procedures, with the corresponding frozen sperm sample.

### Mouse strains and husbandry

Mice were maintained according to Australian Standards for Animal Care under protocol A2018/36 approved by The Australian National University Animal Ethics and Experimentation Committee for this study. The *Zic5*^*tm1Sia*^ targeted null allele (MGI:3574814) of *Zic5* (*Zic5*^*−*^) (Furushima *et al.*, 2005) and the *Zic5*^*Ki*^ ENU allele were backcrossed for 10 generations to the C3H/HeH inbred strain and subsequently for >10 generations to the C57Bl6/J inbred strain. Mice were maintained in a light cycle of 12 hr light: 12 hr dark, the midpoint of the dark cycle being 12 AM. Mice were genotyped by PCR screening of genomic DNA extracted from ear biopsy tissue (Arkell *et al.*, 2001). For the *Zic5*^*tm1Sia*^ strain, genomic DNA (50 ng) was amplified for High Resolution Melt Analysis (HRMA) using IMMOLASE DNA Polymerase using the primers and PCR conditions described for the Z5N assay (Supplementary information; Thomsen *et al.*, 2012). For the *Zic5*^*Ki*^ strain, genomic DNA (50 ng) was amplified for Allelic Discrimination using the following primers and probes: Ark1271, forward (5’ GGC CTT CCT GCG CTA CAT G 3’), Ark1272, reverse (5’ GGT CCA GCC ACT TGC AGA TG 3’), Probe 1 (wild-type; 5’ VIC – TCC CGC TTG ATT GG 3’), Probe 2 (mutation; 5’ FAM – CTC CCG CCT GAT T 3’) and Platinum Quantitative PCR SuperMix-UDG w/ROX (Life Technologies). The products were amplified and analysed on an ABI StepOne PCR machine.

### Whole mount in situ hybridisation

Whole mount in situ hybridisation (WMISH) to *Foxd3* was performed as previously described (Elms *et al.*, 2003; Barratt and Arkell, 2020b, 2020a). A minimum of four embryos for each genotype were compared at eight somites to precisely stage-matched wild-type littermates.

### Statistical analysis

For cell-based assays where one representative experiment is shown, the standard deviation was calculated from three internal repeats using Excel. For analysis of the pooled raw data (from a minimum of three external repeats) of these assays, GenStat software (VSN International) was used to test for normality (W-test) and to perform a two-way ANOVA and Bonferroni multiple comparison test. Post statistical analysis, the values calculated by the ANOVA and the SEM were normalised to the negative control and relative values plotted.

For statistical analysis of the amount of ZIC5 SUMOylated in response to LiCl treatment, GenStat software was used to perform a paired two-sample *t*-test on data from five independent repeats. For analysis of subcellular localisation and co-localisation, GenStat software was used to perform a regression analysis. Post analysis, predicted means and standard error of difference (SED; subcellular localisation) or standard error (SE; for co-localisation studies) was plotted. Mouse breeding data was tested for altered phenotype frequency between genotypes using a G-test of goodness-of-fit.

## Acknowledgements

We thank the following for the gift of plasmids: S. Tejpar; pFLAG-TCF4, S.M. Huang; pSG5-HA-hUBC9, H. McBride; pEYFPC3-SUMO-1, F. Zafra; pXP2-*Apoe*, Y. Sekido; pcDNA3.1/V5-HisA-β-CATENIN, O. Janne; pFLAG-Ubc9-C93S, R. Niedenthal; UFDS plasmids and D. Saunders; V2-ubiquitin, V1-ORF, V2- ORF. We thank S. Aizawa for the *Zic5* null mouse strain. We thank P.l Denny, D. Quwalid and M. Fray for technical assistance with the isolation of the *Zic5* K393R allele. We thank R. Houtemeyer for technical assistance with luciferase reporter assays and helpful discussions. This work was supported by the award of a Sylvia and Charles Viertel Senior Medical Fellowship to RMA.

## Author Contribution statement

Conceptualization: R.G.A., R.M.A.; Methodology: R.G.A., R.M.A.; Validation: R.G.A., R.M.A.; Formal analysis: R.M.A., H.B, J.A; Investigation: R.G.A., H.B., N.W., J.A, K.B and K.N. Resources: R.M.A.; Writing - original draft: R.G.A.; Writing - review & editing: H.B., K.E.M, K.S.B, R.M.A.; Visualization: R.G.A., H.B., J.A, K.B and K.N.; Supervision: R.M.A; Project administration: R.M.A.; Funding acquisition: R.M.A.

## Supplementary Figure legends

**Supplementary Figure 1:**
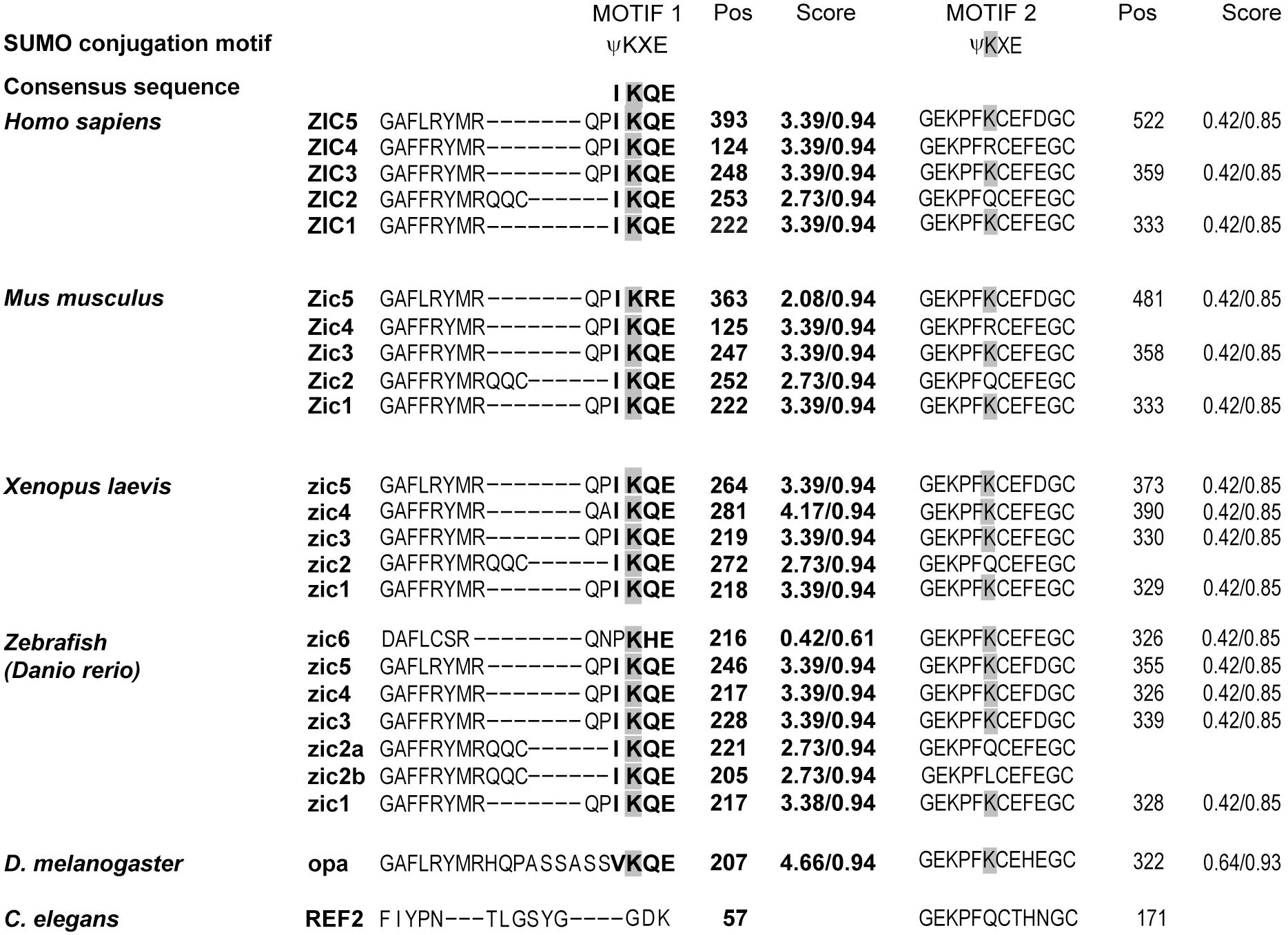
Conservation of SUMO motifs and blots illustrating SUMOylation of ZIC5 within the ZF-NC domain. Conservation of the Motif 1 and Motif 2 consensus SUMOylation sites in Zic proteins from a range of vertebrate species as well as *D. melanogaster* and *C. elegans*. Although most species show a high level of conservation, the consensus SUMOylation sites are missing from *C. elegans*. The first score for each motif is computed by SUMOsp and the second by SUMOplot.

**Supplementary Figure 2:**
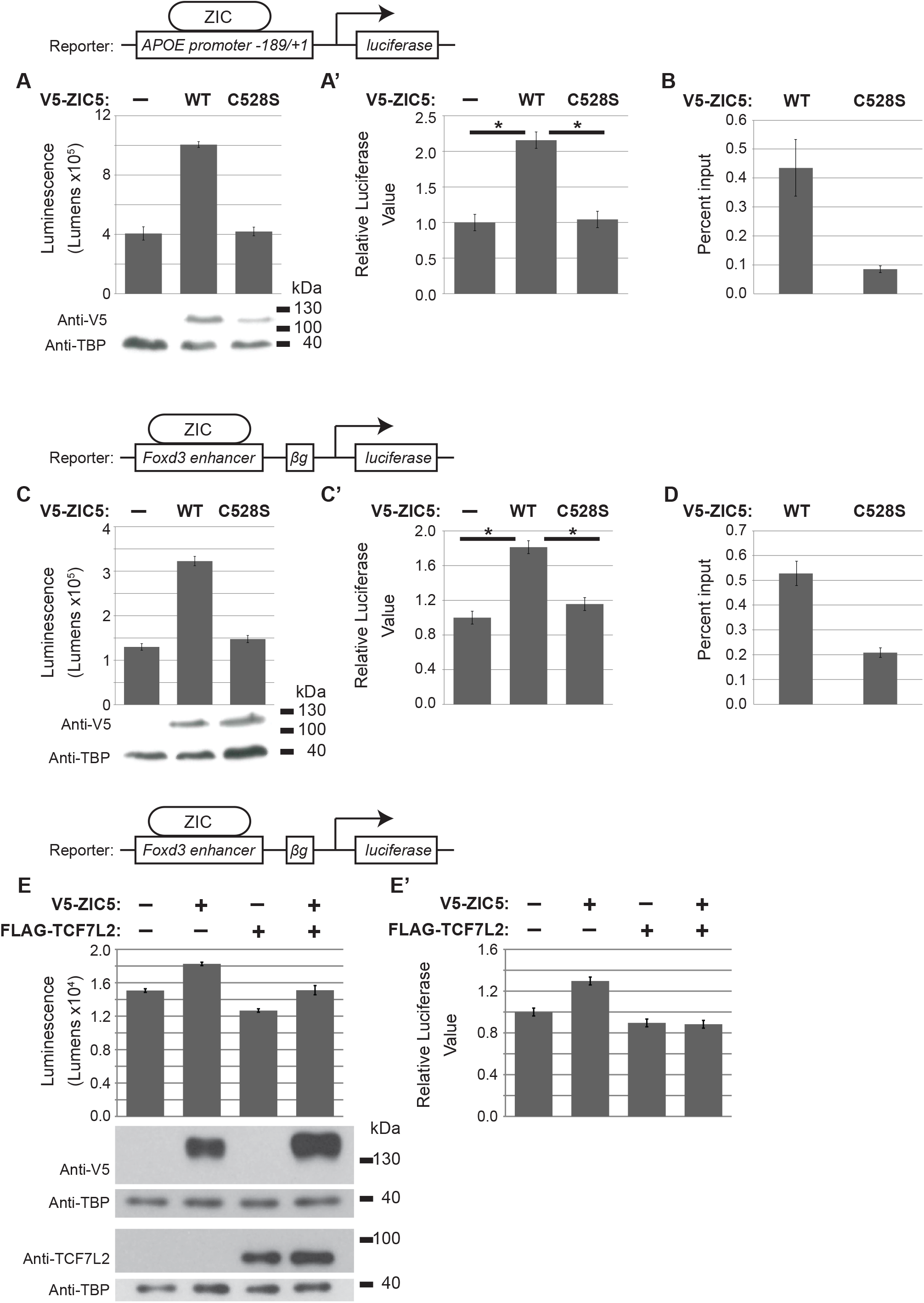
ZIC5 DNA binding is required for activation of *Apoe* and *Foxd3* reporter constructs. (A and A’) Wild-type ZIC5 protein but not the SUMO-incompetent form of ZIC5 (ZIC5-C528S) is able to transactivate the *Apoe* reporter construct. (B) qRT-PCR to *Apoe* promoter fragment demonstrating enrichment of V5-ZIC5-WT but not V5-ZIC5-C528S using plasmid IP. Error bars = SD from three internal repeats. (C and C’) Wild-type ZIC5 protein but not the SUMO-incompetent form of ZIC5 (ZIC5-C528S) is able to trans-activate the *Foxd3* reporter construct following co-transfection into HEK293T cells. (D) qRT-PCR to *Foxd3* promoter fragment demonstrating enrichment of V5-ZIC5-WT but not V5-ZIC5-C528S using plasmid IP. Error bars = SD from three internal repeats. (E and E’) TCF7L2 does not cooperate with ZIC5 to activate Foxd3 expression. Luciferase reporter activity in HEK293T cells to measure ZIC dependent transcription using the *Foxd3* enhancer based reporter construct. Over-expression of TCF7L2 alone or in conjunction with ZIC5 does not activate this reporter. (A, C, E) Raw data and corresponding WB of overexpressed proteins from one representative experiment. Error bars = SD from three internal repeats. (A’, C’, E’) Pooled data from three external repeats (normalized to the background which is set to one). Error bars = SEM (ANOVA), *: *p*<0.01 (A’, C’), *: *p*<0.05 (E’) two-way ANOVA with Bonferroni multiple comparison test.

**Supplementary Figure 3:**
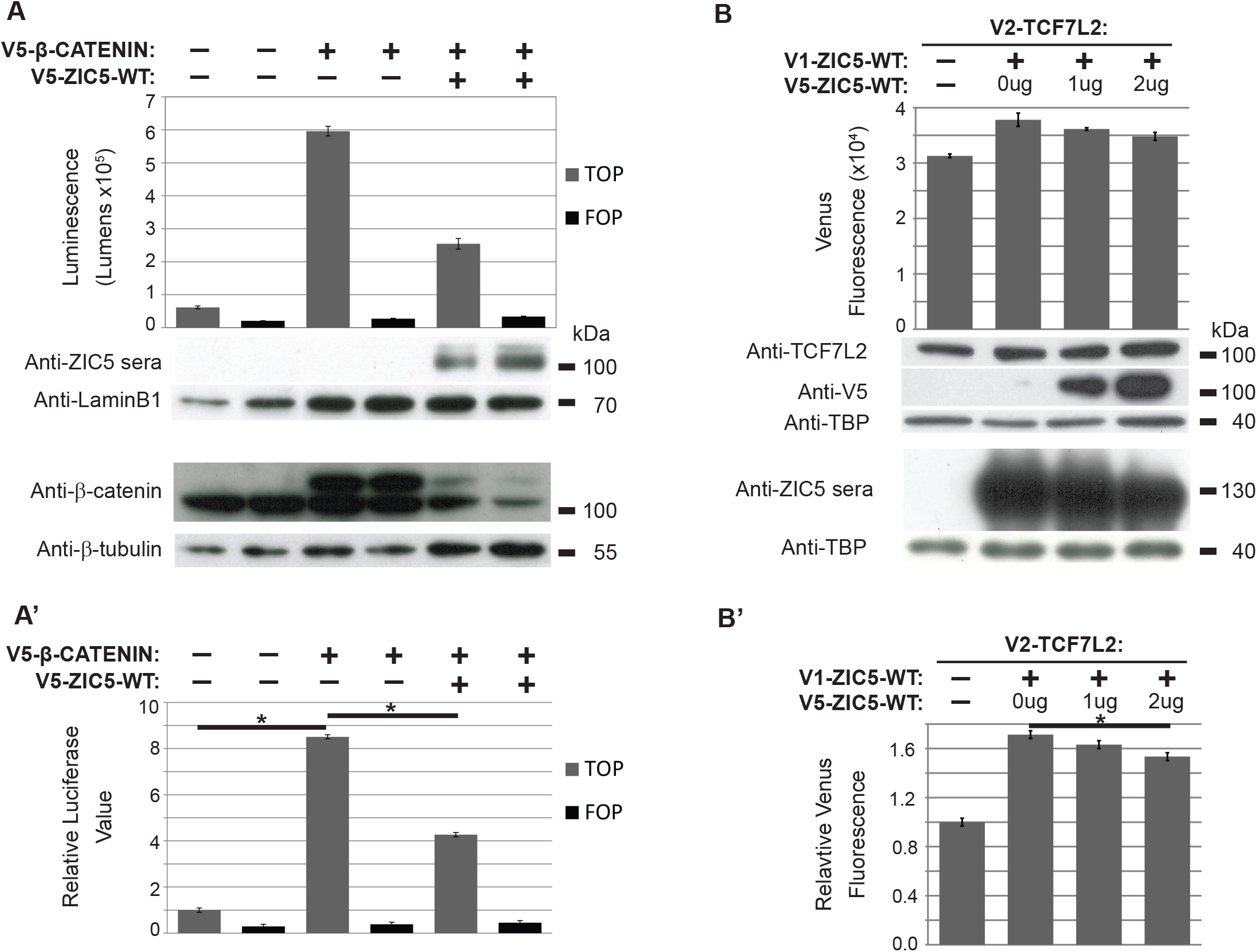
Control assays for Wnt reporter and BiFC assays. (A and A’) Specific stimulation of the TCF reporter construct in the presence of β-catenin. Expression constructs (V5-β-CATENIN and V5-ZIC5-WT) were co-transfected with the TOPflash (TCF) reporter construct (Grey bars) or the FOPflash (mutant TCF) reporter construct (Black bars) into HEK293T cells and luciferase activity subsequently measured. In all cases, little stimulation of the FOPflash construct was observed. (A) Raw data and corresponding WB of overexpressed proteins from one representative experiment. Error bars = SD from three internal repeats. (A’) Transformed data (normalized to the background luciferase value which is set to one) pooled from three external repeats. Error bars = SEM (ANOVA), *: *p*<0.01, two-way ANOVA with Bonferroni multiple comparison test. (B and B’) The presence of non-fluorescent tagged ZIC5 (V5-ZIC5-WT) competes with the split tagged ZIC5 (V1-ZIC5-WT) in the BiFC assay. (B) Raw data and corresponding WB of overexpressed proteins from one representative experiment. Error bars = SD from three internal repeats. (B’) Transformed data (normalized to the background fluorescence value which is set to one) pooled from three external repeats. Error bars = SEM (ANOVA), *: *p*<0.05, two-way ANOVA with Bonferroni multiple comparison test.

**Supplementary Figure 4.**
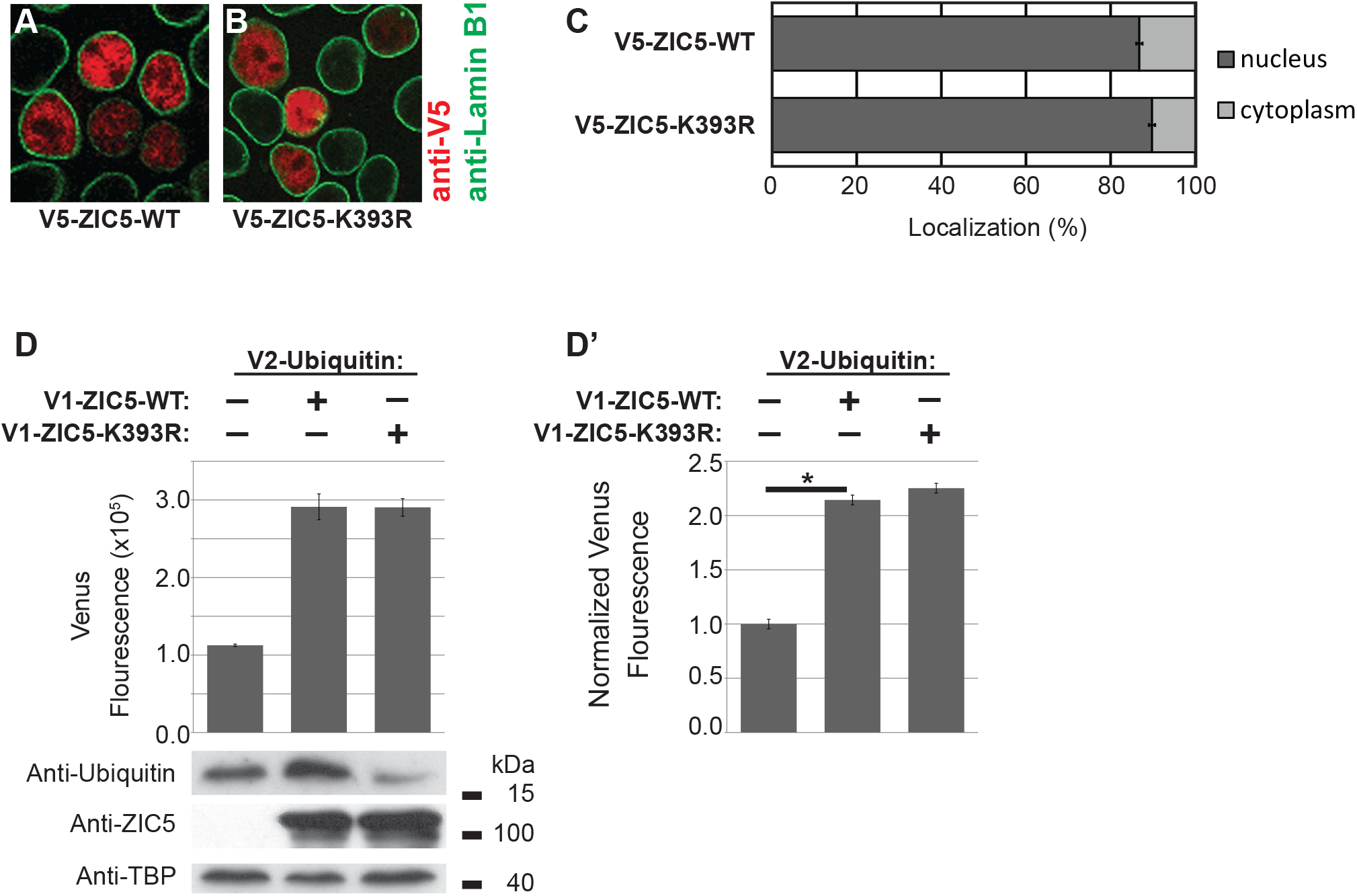
SUMOylation does not alter ZIC5 localization or modification by Ubiquitin. (A-C) SUMOylation does not alter the subcellular localization of ZIC5. The relative distribution of ZIC5 protein was analysed by immunofluorescence microscopy following transfection into HEK293T cells. The ZIC5 protein was identified by hybridization with α-V5 (red) and the nuclear envelope marked by α-Lamin B1 (green). Representative, merged images (shown in A and B) demonstrate the predominately nuclear location of both the WT and K393R forms of ZIC5. (C) The average relative amounts of protein in the nuclear and cytoplasmic compartments calculated from quantification of WT and K393R forms of ZIC5. Graph shows pooled data from three independent experiments (at least 100 cells scored per experiment), Error bars = SED (regression analysis). The two proteins were not found to be significantly different at the *p*<0.01 when compared using regression analysis. (D, D’) ZIC5 is ubiquitinated at a lysine other than 393. BiFC assay between Venus N-terminal (V1) labelled ZIC5 and Venus C-terminal (V2) labelled ubiquitin. The increased fluorescence indicates these proteins interact and do so in the same manner when lysine 393 is mutated, indicating that this lysine is not the modified residue. (D) One representative experiment, Error bars= SD of three internal repeats. (D’) Average values from three external repeats, Error bars = SEM, *: *p*<0.01, two-way ANOVA with Bonferroni multiple comparison test.

**Supplementary Figure 5.**
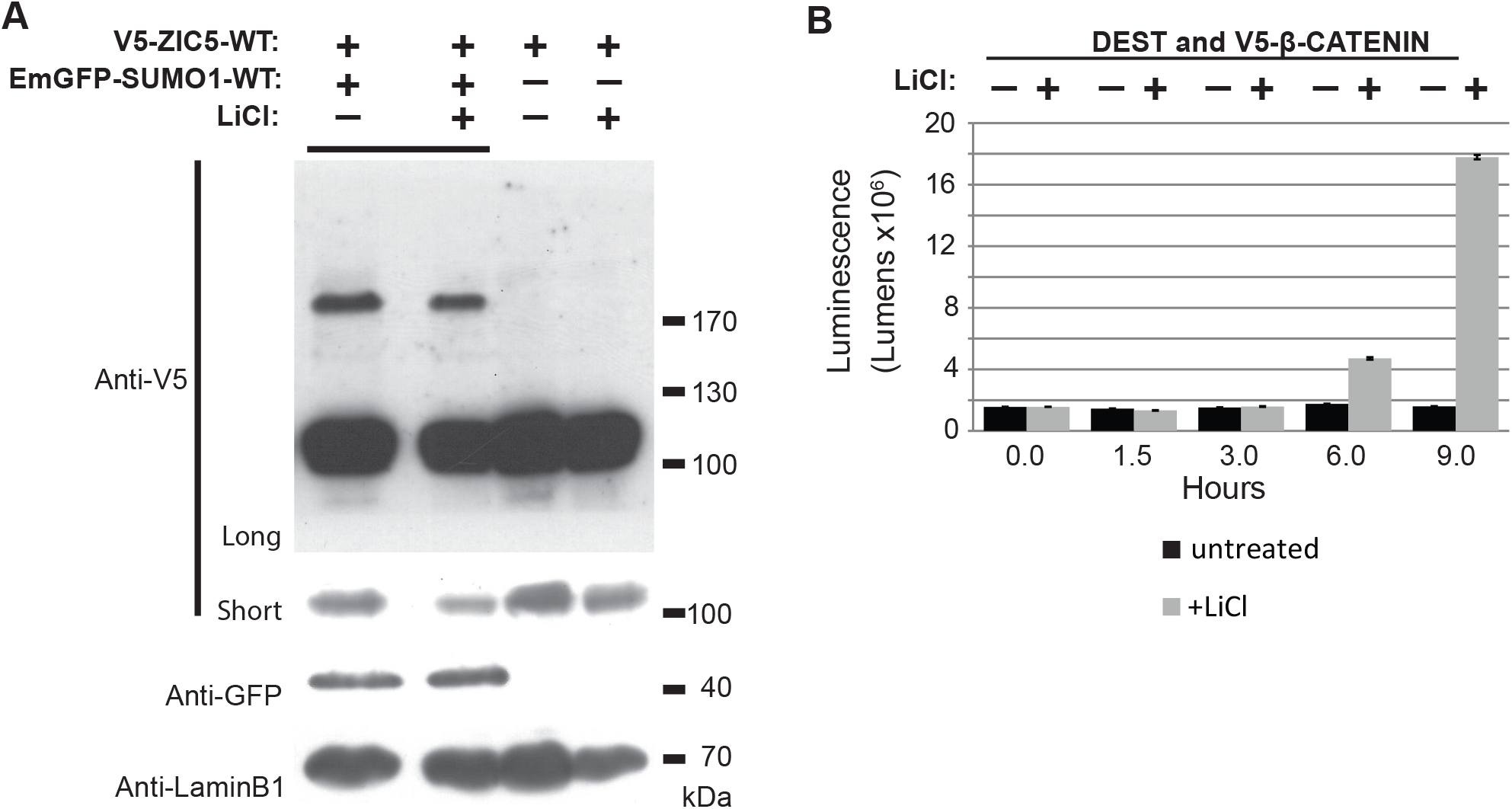
Activation of WNT signalling increases the proportion of SUMOylated ZIC5 and β-catenin mediated transcription. (A) Expanded view of the WB shown in Figure 4A. The lanes shown in Figure 4A are marked by the bar at the top of the WB. The additional two lanes show that the high molecular weight band quantified in the experiment is SUMO1-dependent. (B) The time-course of luciferase production in a TOPflash assay in the presence or absence of LiCl, from one representative experiment. Error bars = SD from two internal repeats. Based on this analysis the 1.5 hour time-point was omitted from the experiments shown in Figure 4B.

